# Allosteric regulation in CRISPR/Cas1-Cas2 protospacer acquisition mediated by DNA in association with Cas2

**DOI:** 10.1101/2020.10.30.361584

**Authors:** Chunhong Long, Liqiang Dai, Chao E, Lin-Tai Da, Jin Yu

## Abstract

Cas1 and Cas2 are highly conserved proteins across CRISPR-Cas systems and play a significant role in protospacer acquisition. Here we study the protospacer (or ps) DNA binding, recognition, and response to cleavage on the protospacer-adjacent-motif complementary sequence or PAMc by Cas1-Cas2, implementing all-atom molecular dynamics simulations. First, we noticed that two active sites of Cas1&1’ bind asymmetrically to two identical PAMc in the simulation. For psDNA containing only one PAMc to be recognized, it is then found that the non-PAMc association site remains destabilized until after the bound PAMc being cleaved. Thus, correlation appears to exist between the two active sites, which can be allosterically mediated by psDNA and Cas2&2’ in bridging. To substantiate such findings, we further simulated Cas1-Cas2 in complex with synthesized psDNA sequences psL and psH, which have been measured with low and high efficiency in acquisition, respectively. Notably, such inter-site correlation becomes largely enhanced for Cas1-Cas2 in complex with psH, and remains low with psL. Hence, our studies demonstrate that PAMc recognition and cleavage in one active site of Cas1-Cas2 allosterically regulates non-PAMc association/reaction in the other site, and such allosteric regulation is mediated by non-catalytic Cas 2 and DNA protospacer in acquisition.

## INTRODUCTION

Bacteria and archaea use clustered regularly interspaced short palindromic repeats (CRISPR) along with associated protein (CRISPR-Cas) adaptive immune systems to protect against invading foreign nucleic acids from phages and plasmids (1-5). The CRISPR-Cas system captures the foreign DNA segments or protospacers and integrate them into the CRISPR array that consists of identical short repeats and variable spacers of similar sizes (1,2,6). The CRISPR-Cas system works in three steps: First, in the protospacer acquisition or adaptation stage, a new spacer is captured and stored in the host genomic CRISPR array (7-9), which is achieved by the highly conserved Cas1-Cas2 protein complex. Second, the CRISPR locus is transcribed and processed into short mature CRISPR RNA (crRNA), which then binds to additional Cas proteins and forms a protein-crRNA complex (2,10). Last, at the targeted interference step, the secondary invading nucleic acid complementary to crRNA is recognized and degraded precisely by the protein-crRNA complex (11,12). Although the overall mechanism of CRISPR-Cas immune system has been well received, some of key steps remain to be elucidated. In particular, the acquisition conducted by Cas1-Cas2 is a least understood step, even though this step is fundamental and ubiquitous to all types of CRISPR systems (9,13-18).

According to the evolutionary classification of the CRISPR-Cas system and Cas genes, Cas1 and Cas2 are the only two Cas proteins conserved across all CRISPR-Cas systems (19-23). Previous biochemical studies identify Cas1 and Cas2 as metal dependent nucleases (24,25). Cas1 is capable of cleaving DNA of various forms in a sequence-independent manner, and demonstrates catalytic activity in the protospacer acquisition. Cas2 is found to be able to cleave single and double-stranded DNA, yet no catalytic activity of Cas2 has been shown in the protospacer acquisition (16,26). Correspondingly, the functional role of Cas2 in the acquisition remains to be elucidated.

The conserved Cas1 and Cas2 proteins can assemble into 2-fold symmetric or dimeric Cas1-Cas2 complex (27,28), with two active sites formed by two Cas1 proteins (Cas1a&1b and Cas1a’&1b’), respectively, while two Cas2 proteins (Cas2&2’) are sandwiched in between the dimeric Cas1. Such a Cas1-Cas2 complex captures the protospacer DNA (or psDNA) before integrating it into the host CRISPR locus. Such an acquisition or adaptation process includes four sub-steps: psDNA binding and selection, the 3’ overhang cleavages, integration, and DNA repair (5,17,18,27,29-31)(see **Figure 1A**). In this study, we focus on the first two sub-steps, i.e., the psDNA binding/selection and cleavages at two active sites. During the psDNA binding, the PAM (protospacer-adjacent motif) recognition is key for the selection (27,28,32)(see **Figure 1B**). It has been shown that PAM is important for recognition and selection of the protospacer during the acquisition (14,15,33). In particular, psDNA flanked by the correct PAM (GAA) can be cleaved and integrated efficiently into the CRISPR array, which ensure that foreign psDNA containing the right PAM (GAA) is incorporated (32,34,35). Meanwhile, for crystal structures captured for the psDNA acquisition complex of Cas1-Cas2 (27), symmetrical PAM complementary sequence or PAMc (CTT) were constructed at the nucleotides 28-30 in the two 3’ overhangs of the DNA protospacer, and both active sites (site 1 and site 2) formed by Cas1a and Cas1a’ are indeed bound with PAMc. Correspondingly, the 3’ PAMc (CTT) sequence in current system is supposed to be cleaved between C and T, and C remains with the spacer to be integrated to the CRISPY array (see the schematics in **Figure 1A** *bottom*).

**Figure 1.**
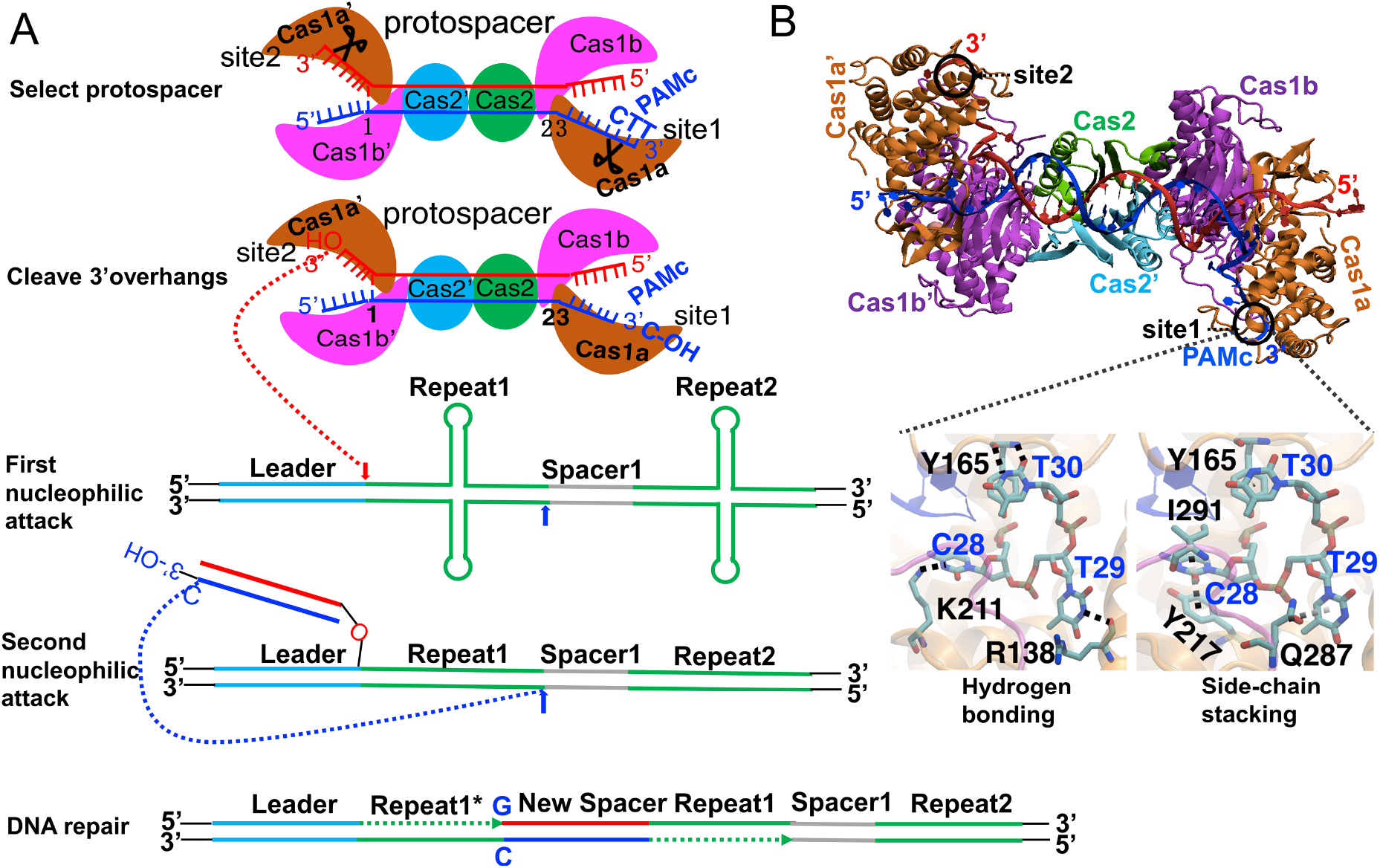
**(A)** Schematics of protospacer (ps) DNA acquisition or adaption into the host CRISPR array for the CRISPR-Cas adaptive immunity. The acquisition process includes four sub-steps (from *top* to *bottom*): protospacer binding and selection, 3’ overhang cleavage, integration, and DNA synthesis/repair. During the integration step, PAMc sequence C’-OH (cleavage at site1 of Cas1-Cas2) is integrated into the Spacer side of Repeat 1 in the CRISPR array (likely via the second nucleophilic attack), while non-PAMc (cleavage at site2) is integrated into the Leader side of Repeat 1 (likely via the first nucleophilic attack) (18,27). **(B)** Crystal structure of *E.coli* Cas1-Cas2 bound to dual-forked psDNA (PDB:5DQZ) (27). Cas1a and Cas1a’ are colored in orange. The Cas1b and Cas1b’ are colored in magenta. Cas2 and Cas2’ are shown in cyan and green, respectively. The two active site1 and site2 are shown with black circles. In the crystal structure, both sites are bound with PAMc (CTT). The PAMc recognition via hydrogen bonding (HB) and amino-acid side-chain stacking in the binding pocket are indicated in black and gray dotted lines (*bottom left* and *right*), respectively.

Upon the psDNA association and the 3’overhang PAMc recognition, the CTT can be cleaved by the Cas1-Cas2 complex at one active site, due to high specificity of such reaction (17,18,27,28). Meanwhile, the other active site, which generally binds to a non-PAMc at the other 3’overhang, would also conduct the catalytic cleavage, though it is not clear whether the cleavage can proceed sufficiently fast on the non-PAMc. Notably, high-resolution structural studies have caught a half-site intermediate integration complex of Cas1-Cas2 (29), which has its non-PAMc side 3’-OH linking to the CRISPR between the leader and repeat1, as if it happens as the first nucleophilic attack on the CRISPR array (see **Figure 1A** schematics). If the non-PAMc cleavage conducted by Cas1-Cas2 happens much slower than the first PAMc cleavage, then it is likely that an alternative half-site intermediate integration complex emerges, which has its PAMc 3’-OH linked to the CRISPR as the first nucleophilic attack (i.e. between the repeat1 and the previous spacer or spacer1). The existence of the first half-site integration complex of Cas1-Cas2, with the non-PAMc 3’overhang linked to CRSIPR, indeed suggests that the non-PAMc needs to be cleaved (at site2, labeled for convenience) almost as fast or immediately after the PAMc cleavage (at site1). We therefore want to understand how the non-PAMc site can be cleaved sufficiently fast in the Cas1-Cas2 acquisition complex. We hypothesized that some communication exists between the two active sites in the Cas1-Cas2 complex so that the efficient non-PAMc cleavage is enabled.

We accordingly conducted atomistic molecular dynamics (MD) simulations on the high-resolution crystal structure of the Cas1-Cas2 complex in association with a dual-forked psDNA (27,28), which contains two PAMc on the two fork regions, respectively. In this dual PAMc complex, the psDNA lies on the surface of Cas1-Cas2 in a head-to-head orientation (see **Figure 1B**). Two identical PAMc are bound by the two active sites (site 1 and site 2), respectively. We simulated the original Cas1-Cas2 binding complex with the dual PAMc, and identified instead an asymmetrical equilibrium binding pattern between the two active sites. Then we modified one PAMc to a non-PAMc (at site 2), while kept the other PAMc intact (at the site 1), so that to mimic PAMc binding and recognition in a general condition. We subsequently examined the Cas1-Cas2 systems from the psDNA binding to pre-catalytic and to a half-site post-catalytic state (right after the first or the PAMc cleavage), and monitored structural and correlation dynamics between two active sites in respective systems. To substantiate the initial findings upon the hypothesis, two synthetical psDNA sequences, which were experimentally identified with low and high acquisition frequencies or efficiencies (36), were introduced into the Cas1-Cas2 psDNA simulation systems. Via comparative studies, we verified negative cooperativity between the two active sites, which relies on allosteric propagation from the site1-PAMc to the site2-non-PAMc, as being mediated by the psDNA in acquisition and non-catalytic Cas2&2’ in close association with the psDNA, sandwiched well between Cas1a&1b (with site1) and Cas1a’&1b’ (with site 2).

## MATERIAL AND METHODS

We illustrate how we constructed the Cas1-Cas2 psDNA binding structure, the pre-catalytic and the half-site post-catalytic structures, along with MD simulation setup. The original psDNA was then modified into sequences with high/low acquisition efficiency (psH/psL), according to synthetic experimental construct (36). In addition, we explain how dynamic correlations are measured between the site1-PAMc and the rest part of the protein-DNA complex from the simulations.

### Constructing Cas1-Cas2-psDNA structural complexes at various states, with dual or single PAMc on the psDNA

The high-resolution structure of Cas1-Cas2 in complex with a dual-forked psDNA containing two identical PAMc (CTT) was obtained from Wang Lab (PDB:5DQZ) (27). The 2-fold symmetric structure contains four Cas1 and two Cas2 polypeptides, with the Cas2 dimer (Cas2&2’) being sandwiched between the two Cas1 dimers (Cas1a&1b and Cas1a’&1b’). The psDNA contains a 23-bp duplex in the middle, which binds onto the flat surface provided primarily by Cas2&2’. The two 3’ overhangs at the forked regions of psDNA containing two PAMc thread into the binding sites formed by Cas1a&1a’, termed site1 and site2, respectively, with residue 278 to 305 of Cas1b&1b’ covering the catalytic pocket. In the original psDNA binding complex of Cas1-Cas2, there are two magnesium ions in each of the Cas1b and Cas1b’ non-catalytic pockets (see Supplementary **Figure S1A** *top*). Next, by using another crystal structure from Doudna Lab (PDB: 5DS5) (28), we inserted two additional magnesium ions into both of the Cas1a and Cas1a’ catalytic pockets (i.e., active site1 and site2, Supplementary **Figure S1A** *bottom*), which are indispensable for catalysis, so that we constructed the pre-catalytic state structure. Equilibrium MD simulations (200 ns) were conducted for both the binding and pre-catalytic structures. The RMSDs for both structures during the equilibrium simulation are provided (in Supplementary **Figure S2**).

Besides the original binding complex with psDNA containing two PAMc, we constructed systems with psDNA containing only one PAMc and one non-PAMc, starting from the two-PAMc crystal structure. In particular, we focused on the systems with site1 bound PAMc (CTT) and site2 associated with non-PAMc (TTT). Additionally, we also constructed a binding complex with site1 associated with non-PAMc and site2 bound PAMc, as well as a binding complex with both site1 and site2 in association with non-PAMc. All these psDNA binding complexes of Cas1-Cas2 were all subjected to 200-ns equilibrium MD simulations.

To further construct a half-site post-catalytic intermediate complex for the system with single PAMc (i.e., site1-PAMc and site2-non-PAMc), we modified the original crystal structure: the PAMc (CTT) bound with site1 was cleaved (as in the post-catalytic state, see Supplementary **Figure S1B**), in which the TT sequences of PAMc were cut and then removed, as being catalyzed via an endonuclease reaction, while the site2 containing the non-PAMc (TTT) remained intact (as in the pre-catalytic status). The half-site post-catalytic intermediate complex was also subjected to a 200-ns equilibrium MD simulation (with RMSD shown in Supplementary **Figure S2**).

### Set up for performing MD simulations

To conduct the MD simulation, the Cas1-Cas2 in complex with the psDNA was solvated with explicit TIP3P water in a cubic box with a size of 180 Å (with a total of ~182,500 water molecules, and a minimum distance between the protein complex and the boundary of the box at 18 Å) (37). As being pointed out in previous MD studies (38), it is essential to make the water box sufficiently large in stabilizing the current system. Initially, we implemented a water box with a size of 150 Å (containing ~ 100,600 water molecules, with the minimum distance between the protein complex and the boundary of the box at 15 Å) in simulating the original binding complex. However, neither of the active sites (site1 and site2) could be stabilized by the PAMc in association. The increasing the box size to 180 Å then stabilizes one PAMc binding site (see Supplementary **Figure S3**).

The protonation status of His195 and His208 in the binding pocket were set as HID in order to allow stable association with magnesium ions, since nitrogen in epsilon position (Nε) is close to the positively charged Mg^2+^ so that the protonation is not preferred. Note that the protonation status of histidine residue includes HID, HIE and HIP, corresponding to Nδ, Nε, and both nitrogen protonated, respectively. The 166 water molecules from the crystal structure (PDB: 5DQZ) (27)were retained. We neutralized the system with counter ions (~573 *Na^+^* and ~527 *Cl*^-^) to make an ionic strength at ~ 0.15M. The full simulation system contained ~570,000 atoms.

Then all MD simulations were performed using GROMACS-5.1.2 package (39,40), and the Amber99sb-ILDN force field with ParmBsc0 nucleic acid parameters were used (41-44). For all simulations, the cutoff of van der Waals (vdW) and the short-range electrostatic interactions were set to 9 and 10 Å, respectively. Particle mesh Ewald (PME) method was used to evaluate the long-range electrostatic interactions (45,46). All the MD simulations were conducted at 1 bar and 310 K, using the Parrinello-Rahman Barostat and the velocity rescaling thermostat (47-49), respectively. Before running the equilibrium simulation, we performed energy minimization on the initial structure for 50,000 steps with the steepest-descent algorithm, followed by 100 ps NVT MD simulation. The time-step was 2 fs and the neighbor list was updated every 5 steps. Position restraints on the heavy atoms of the protein and nucleic acids chains were imposed in the 100-ps NVT simulation, and in the first 10-ns NPT MD simulation at the beginning of each 200-ns equilibrium simulation. Following the constrained simulation, unconstrained MD simulation was still carried out under the same NPT ensemble, with a time step of 2 fs.

### The protospacer DNA (psDNA) sequences and modification to psL/psH

According to the low/high acquisition efficiency of the psDNA sequences (psL/psH) synthesized for the Cas1-Cas2 acquisition experimentally (36), we modified the original psDNA sequences contained in the Cas1-Cas2 complex (from the crystal structure) (27)to the psH/psL sequences, using the w3DNA software (50). The original, psL and psH DNA sequences are listed below.

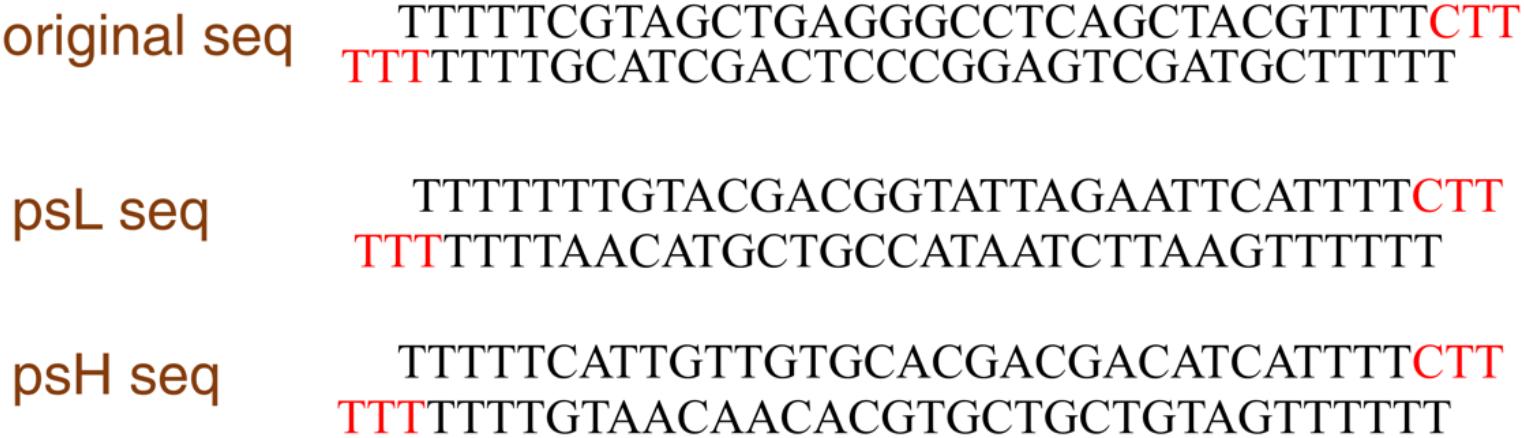

### Correlation calculation from the equilibrium MD simulation

The protein-DNA internal dynamic correlations with the active site1 (bound with PAMc) were calculated to reveal how each residue from the rest of the protein-DNA complex couples with the active site1, via atomic motions sampled from the equilibrium MD simulations. In order to obtain the individual residue correlations with the active site1, we calculated first the pairwise correlations for all amino acids and nucleic acids residues. The correlation between each pair of residues is given by 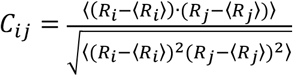, where *R_i_* and *R_j_* are the position vectors of residue *i* and *j*, taking at *C_α_* atom of an amino acid or the center of mass (COM) of the nitrogen atoms (N1 to N4, N6, N7, N9) in the base of a nucleotide and the backbone P atom; 〈…〉 represents averaging over the simulation trajectory (51,52). Accordingly, we obtained a pairwise *N x N* correlation matrix, with *N* =1624 counting all residues from protein and DNA, by averaging over 200 snapshots from each of the 200-ns equilibrium simulation. Finally, we calculated the correlation between residue *i* and the active site1 (without including PAMc) as 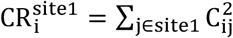, where residue *i* are counted for all C_α_ atoms from protein and the COM of nitrogen and phosphorous atoms in the psDNA.

## RESULTS

By employing all-atom MD simulations, we analyzed the Cas1-Cas2 protospacer binding and selection, via examining formation of potential HBs and base stacking characteristics at the active site. We first examined the original psDNA binding complex with two identical PAMc bound at both sites (site1 and site2); then we focused on the modified system, with one PAMc (CTT) bound at site1 and one non-PAMc (TTT) in association with site2. For such one-PAMc system, we examined not only the psDNA binding state, but also a pre-catalytic state (with catalytic magnesium ions bound to the active sites), and a half-site post-catalytic state (with PAMc cleaved at site1, as being catalyzed via an endonuclease reaction). For each state of the one-PAMc system, the protein-DNA internal dynamics correlations with the active site1 are calculated to reveal allosteric communication from the PAMc-binding site to the non-PAMc binding site or the rest part of the Cas1-Cas2 psDNA complex. Cas1-Cas2 in complex with the synthetic psL and psH DNA sequences are presented further to demonstrate the modulated allosteric effects and the role of psDNA and Cas2 in the communication.

### Cas1-Cas2 bound with two identical PAMc are asymmetrically stabilized at one site and nonstabilized at the other site

We performed a 200-ns equilibrium MD simulation to the original psDNA binding complex of Cas1-Cas2, which was crystalized with 2-fold symmetry, and with two identical PAMc bound symmetrically at both active site1 and 2 (PDB:5DQZ) (27)(see Supplementary **Figure S4**). According to the HBs and stacking interactions formed between protein and PAMc at site1 and site2 in the crystal structure, we examined whether these interaction characteristics maintain in the equilibrium simulation. In order to keep simulation model close to the crystal structure, positional restrains were implemented for up to 10-ns at the beginning of the simulation (see Methods). When simulated in a sufficiently large water box (with a distance between the protein complex and boundary ~20 Å, and the simulation system ~0.5 million atoms), the HBs and stacking interactions at site1 were maintained well during the simulation (see **Figure 2A-C**). In contrast, at the site2, we found that the expected HB between residue R138 and T29 of PAMc (CTT) was broken at ~ 70 ns of the simulation, and the distance between them reached up to ~12 Å (**Figure 2B**). The expected stacking between Q287 and T29 at binding site2 also became unstable after ~ 100 ns, as the distance between the centers of mass (COMs) of residue Q287 and T29 reached above ~6 Å (**Figure 2D**). The binding configurations of the active site1 and site2 at the end of the simulation are shown in **Figure 2E** and **F**, respectively. The observations indicate that the two active sites maintain asymmetrical binding patterns to PAMc, even though the crystal structure shows twofold symmetry and with two PAMc association with site1 and site2 equivalently. The asymmetrical PAMc binding dynamics of the two active sites of Cas1-Cas2 also implies that it can easily accommodate binding/recognition to psDNA containing only one PAMc, considering that two PAMc separated by ~30 bp is a rare configuration to be found on psDNA. Below we focus on the system that is commonly incurred, i.e., Cas1-Cas2 binding to psDNA with only one-PAMc.

**Figure 2.**
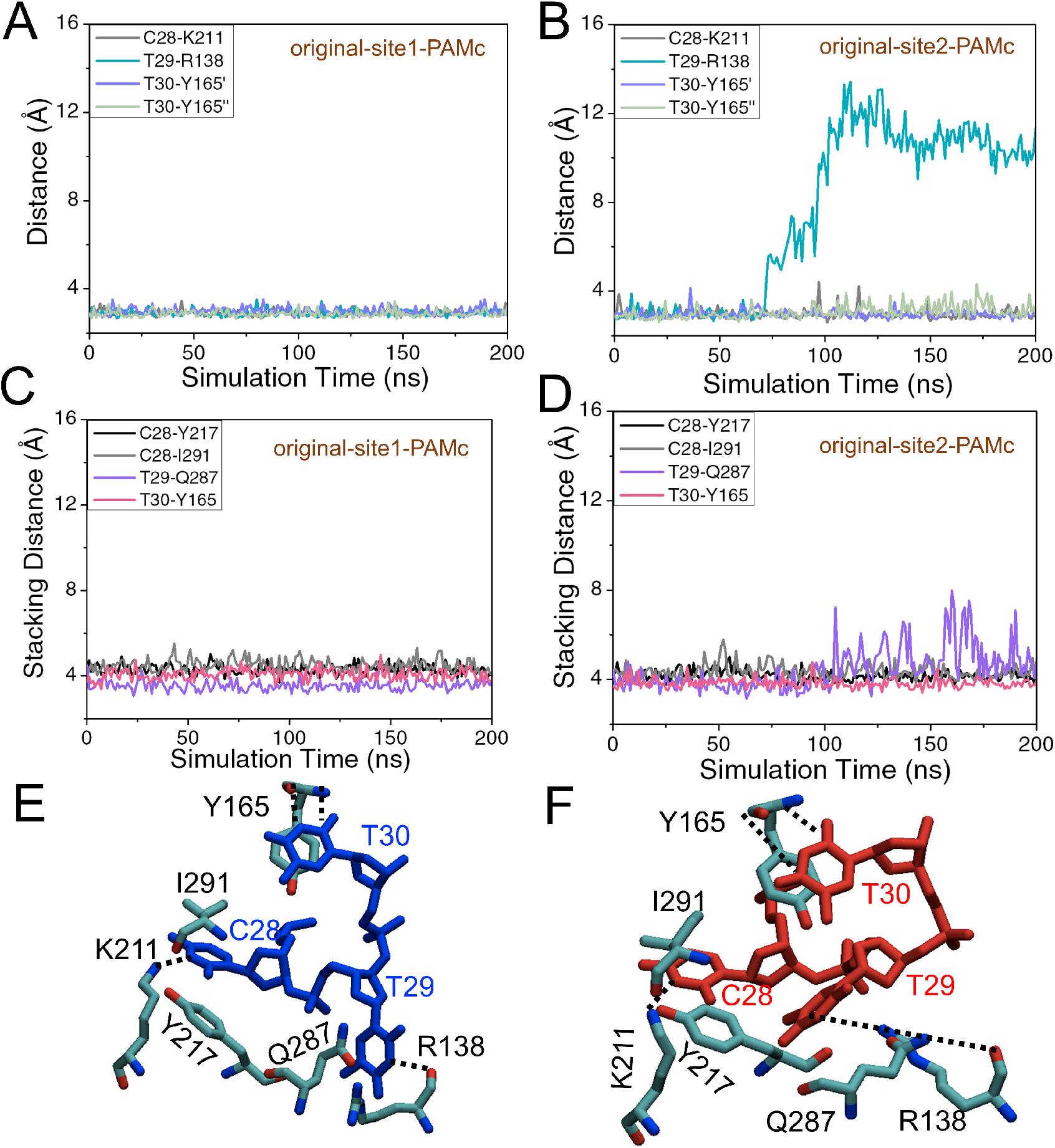
The equilibrium MD simulation of the Cas1-Cas2 complex with dual PAMc psDNA and respective association characteristics at both site1 and site2. **(A)** and **(B)** The expected HB interactions with distances measured between residues R138, Y165, K211 and PAMc (CTT) at binding site1 and site2, respectively. **(C)** and **(D)** The expected stacking interactions with distances measured between the COMs of residues Y165, Y217, Q287, I291 and PAMc at binding site1 and site2, respectively. **(E)** and **(F)** The structural views at site1 and site2, at the end of the 200-ns equilibrium simulations. The PAMc nucleotides are colored blue and red at site1 (E) and site2 (F), respectively. The amino acids are colored by atom types (cyan, red, and blue for carbon, oxygen, and nitrogen, respectively). The expected HBs (according to the crystal structure; may not form in the simulation, e.g. at site2) are highlighted by dotted lines.

### The site2-non-PAMc cannot be stabilized until after the PAMc cleavage conducted at the site1

Except for the original dual PAMc psDNA binding complex of Cas1-Cas2, the following systems were all modified to: site1-PAMc (CTT) and site2-non-PAMc (TTT), and we modeled the system from the psDNA binding to the pre-catalytic state and then to the half-site post-catalytic state (see Methods). Similarly, we checked the expected protein-DNA HBs and stacking characteristics as such for the stabilized PAMc binding and recognition. First, we found that the HB between residue R138 and T29 from non-PAMc (TTT) at site2 was entirely absent in both the binding and pre-catalytic states during respective simulations (see **Figure 3A** and **B**). Second, the expected HB between residue K211 and T28 from TTT at site2 in the pre-catalytic state was also absent (**Figure 3B**). In addition, the stacking interaction between Q287 and T29 at site2 was also unstable, in both the binding and the pre-catalytic states (see Supplementary **Figure S5A** and **B**). Meanwhile, such expected HBs and stacking interactions were stably maintained at site1 bound with PAMc. Hence, the asymmetrical binding configuration of site 1 and site2 does fit well with the PAMc and non-PAMc association, respectively.

**Figure 3.**
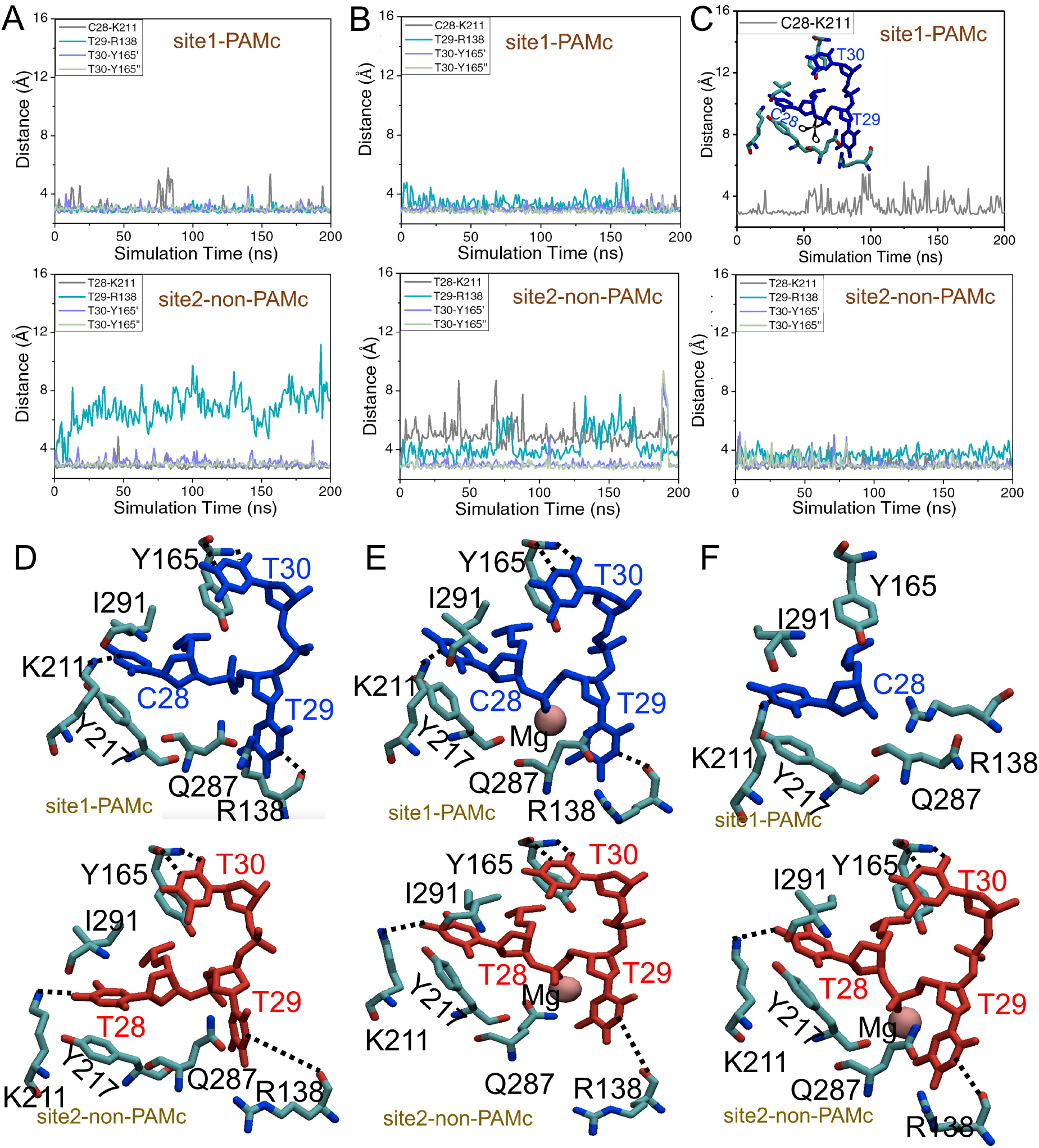
The association patterns at the site1-PAMc and site2-non-PAMc in the equilibrium MD simulation of the Cas1-Cas2 psDNA complexes from the binding to pre-catalytic and to half-site post-catalytic state. The expected HBs with distances measured between residues Y165, R138, K211 at site1-PAMc (CTT; *top*), and similarly at site2-non-PAMc (TTT; *bottom*), in the binding state **(A)**, the pre-catalytic state **(B)**, and the half-site post-catalytic state with PAMc cleaved at site1 **(C)**. The molecular view at the end of the equilibrium MD simulation for the site1-PAMc (*top*) and site2-non-PAMc (*bottom*), from the binding **(D)** to pre-catalytic **(E)** and to half-site post-catalytic state **(F)**. The coloring scheme is the same as Figure 2E and F.

Remarkably, for the half-site post-catalytic state, i.e., after the PAMc was cleaved artificially at site1, the site2 in association with non-PAMc became stabilized, according to the same HBs and stacking characteristics expected for the stable binding configuration (see **Figure 3C** and Supplementary **Figure S5C**). The molecular views toward the end of each 200-ns equilibrium MD simulation, from the binding to the pre-catalytic state and to the post-catalytic state, for both the site1-PAMc and site2-non-PAMc are also provided (see **Figure 3D to F**). From the molecular views, one can directly see that the expected HB between residue R138 and T29 from TTT at site2, for example, which was absent in both the binding and pre-catalytic state system, became present in the half-site post-catalytic state, upon the PAMc cleavage at site1.

These results consequently suggest that the two active sites exhibit negative cooperativity with each other, and such cooperativity only allows one active site to be stabilized in close association with the psDNA for binding and recognition. Consequently, site2 can bind tightly with the non-PAMc only after the PAMc is cleaved at site1, as the close association between site1 and psDNA becomes lost. We thus infer that the cleavage to the PAMc at the corresponding binding/recognition site of Cas1-Cas2 is necessary for the non-PAMc to be bound favorably by the other site and cleaved sufficiently fast.

Besides, we have also examined another two Cas1-Cas2 psDNA complex systems: (i) with site1-non-PAMc (TTT) and site2-PAMc (CTT), and (ii) with both site1-non-PAMc (TTT) and site2-non-PAMc (TTT) (see Supplementary **Figure S6**). For the system (i), we found that site2 with PAMc became stabilized, while site1 with non-PAMc was unstable. For the system (ii), both site1 and site2 became highly unstable with the non-PAMc, e.g., the expected HBs between residue K211 and T28 and between residue R138 and T29 were absent.

### The allosteric propagation from the site1-PAMc to the site2-non-PAMc upon the site1-PAMc cleavage

For each 200-ns equilibrium simulation of the Cas1-Cas2-psDNA complex, from the binding to pre-catalytic and to half-site post-catalytic state, the dynamics correlation within the protein was calculated (see **Methods**), in particular, between the active site1-PAMc and the rest part of the protein-DNA complex (see **Figure 4A**). Note that high self-correlation values (>30) show for Cas1a which hosts the active site1, and for the PAMc regions in close association with the active site1. Residue 93 to 108, residue 200 to 220, and residue 275 to 295 in Cas1b around the active site1 also show high correlations (~ 20) with the active site1, due to local interactions. For non-local regions (Cas1a’&1b’, the rest part of DNA aside from PAMc, and Cas2&2’, labeled in **Figure 4A**), the site2-non-PAMc, particularly located in Cas1a’, demonstrates non-trivial correlations with the active site1 (i.e., up to 5-10 into the post-catalytic state, see **Figure 4B**); psDNA and Cas2&2’ also show slightly increased correlations with the site1 into the post-catalytic state (**Figure 4C** and **D**).

**Figure 4.**
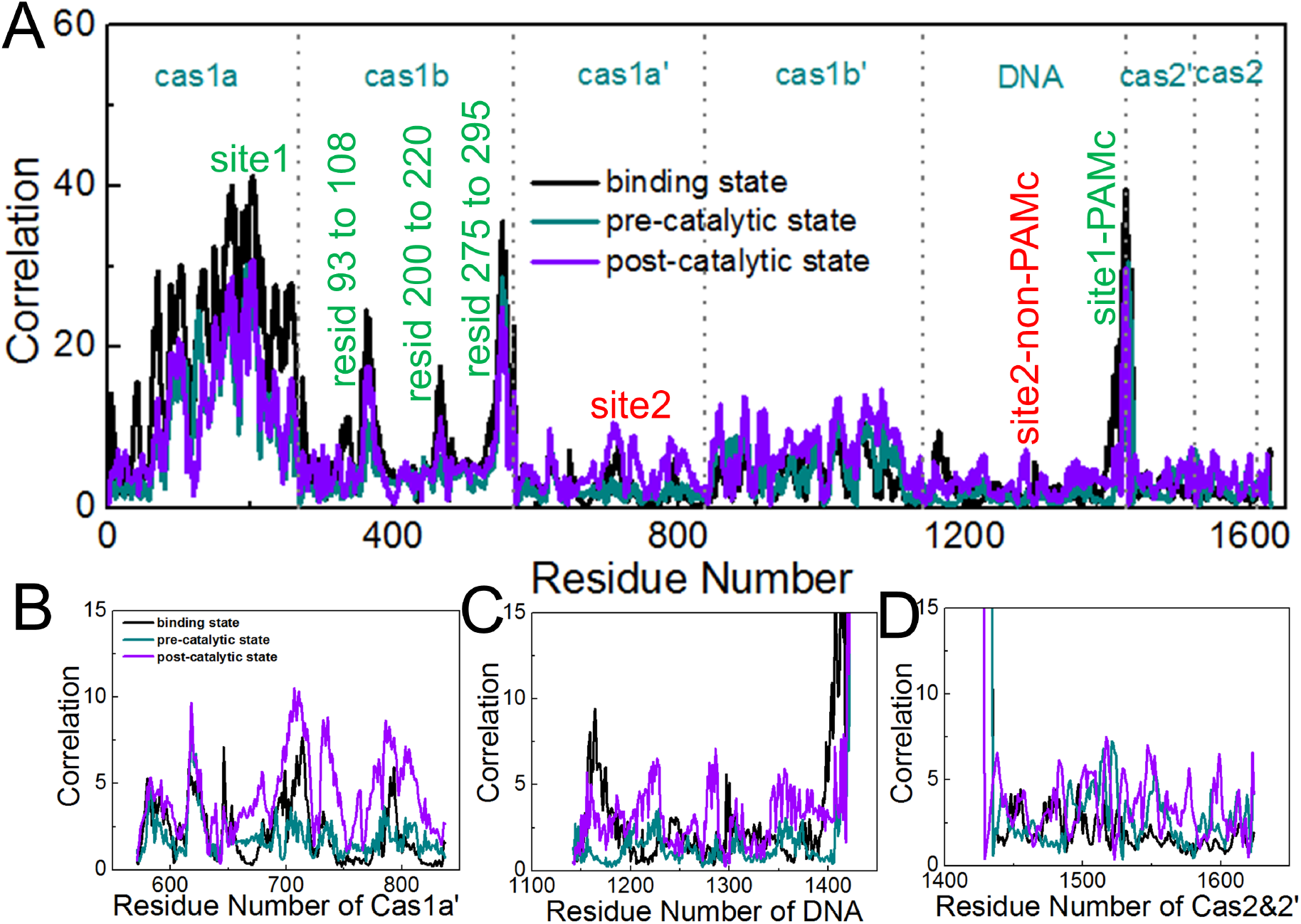
The internal correlation strength between the active site1 and the rest parts of the Cas1-Cas2 psDNA complex (site1-PAMc and site2-non-PAMc) during the equilibrium MD simulations. **(A)** The correlations with the site1 (PAMc) with the rest part, from the binding state (dark line) to pre-catalytic state (cyan) and to half-site post-catalytic state (purple). The residues local around site1 with comparatively high correlations are labeled (green), while remote regions are highlighted (red). **(B-D)** The zoomed-in views of (A) for the correlation patterns between the active site1 and non-local residues from Cas1a’ hosting site 2 (B), psDNA (C), and Cas2&2’ (D).

In comparison with the binding and pre-catalytic state, the protein-DNA complex upon the site1-PAMc cleavage (i.e., at the half-site post-catalytic state) shows lowered self or local correlations with the site1-PAMc, but enhanced non-local correlations with the site1 at remote regions, including Cas1a’&1b’ (hosting site2), psDNA (aside from PAMc), and Cas2&2’. Hence, one sees that the correlation between the two far-apart active site1 (at Cas1a&1b) and site2 (Cas1a’&1b’) exits and strengthens upon the PAMc cleavage at site1. It can be inferred that the lowered local correlation and increased non-local correlation with the active site1 into the post-catalytic state are caused by allosteric propagation of the site1-PAMc cleavage dynamics to the rest part of the protein complex, mediated by psDNA and Cas2&2’, and reaching remotely to the site2-non-PAMc (~ 10.3 nm apart from site1).

For further comparison, we also examined the protein internal correlations in the original psDNA binding complex of Cas1-Cas2 with dual PAMc, i.e., with site1 stabilized and site2 destabilized in the simulation. The correlations were calculated for both the active site1 and site2, as to see how the two sites respectively couple with the rest part of the protein-DNA complex. Overall, the correlations in the dual PAMc system appear larger than that in the single PAMc case. Interestingly, one finds that the remote correlations to the stabilized site1 and to the non-stabilized site2 are similarly large, though the local or self-correlation near the non-stabilized site2 appears weaker than that near the stabilized site1 (see SI Fig S7). Hence, allosteric propagation between the two active sites remote to each other on Cas1-Cas2 appear mutual and comparatively robust, regardless of the local site stability.

### Cas1-Cas2 in complex with psDNA of high acquisition efficiency (psH) shows the most prominent allosteric propagation upon the PAMc cleavage

According to synthetic approach to sequence-dependent protospacer acquisition of CRISPR/Cas1-Cas2, different psDNA sequences can lead to different acquisition frequencies or efficiency (36). In particular, two representative protospacer sequences with particularly low and high acquisition efficiencies identified experimentally were incorporated into our simulation systems (psL/psH), so that we could examine whether allosteric propagation between two active sites persists in the Cas1-Cas2 complex with psL or psH, in comparison with the original psDNA.

Similarly, we performed a series of equilibrium simulations for both the psL and psH binding/pre-catalytic/post-catalytic systems of Cas1-Cas2, and calculated the protein internal correlations between the active site1 (with PAMc) and the rest part of the protein-DNA. Comparing the three systems with the original psDNA, psL, and psH, we found that the overall protein internal correlation is actually highest for the Cas1-Cas2 bound with the psH DNA sequences, in particular, in the half-site post-catalytic state (see **Figure 5A**). The zoomed-in views show more closely the site1 correlation with the Cas1a’ (containing the site2), psDNA, and Cas2&2’ regions (see **Figure 5B to D**). The results thus suggest that the high acquisition efficiency of the psH system can be explained by significant correlations induced inside Cas1-Cas2 and between the two active sites. The results also support the idea that the allosteric propagation from the site1-PAMc to the site2-non-PAMc is regulated by the psDNA and Cas2&2’, which is in close association with the duplex region of the psDNA.

**Figure 5.**
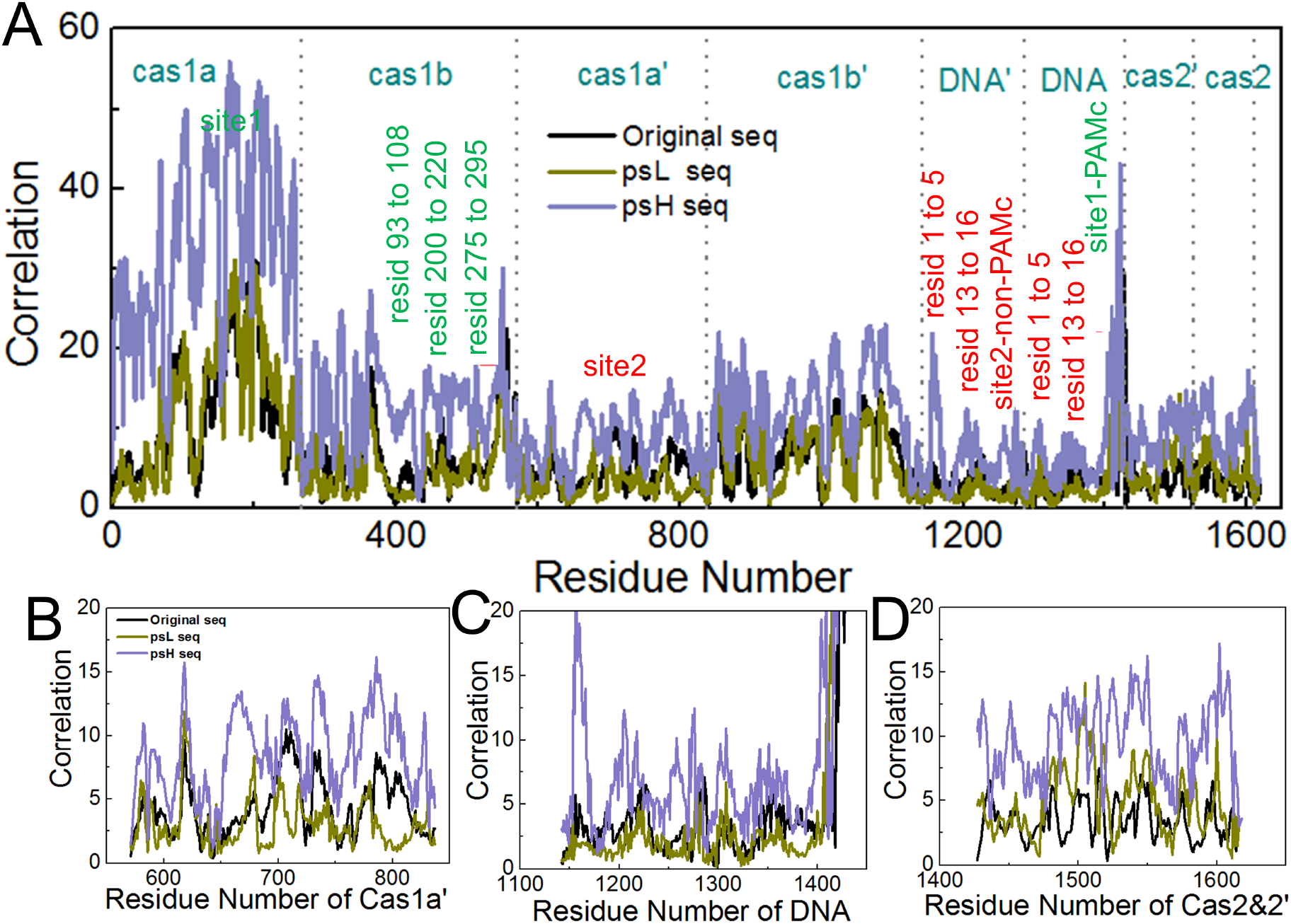
The protein internal correlation from the equilibrium MD simulations between the active site1-PAMc and the rest parts of the Cas1-Cas2 psDNA complex in the half-site post-catalytic state with different psDNA. **(A)** The correlations with the site1 for the original psDNA (black line), psL (ginger), and psH (purple). The residues local around site1 with comparatively high correlations are labeled (green), while remote regions are highlighted (red). **(B-D)** The zoomed-in views of (A) for the correlation patterns between the active site1 and residues from Cas1a’ hosting the site2 (B), psDNA (original/psL/psH) (C), and Cas2&2’ (D).

A color map of the correlation strength (blue/white/red: high/medium/low correlation) on Cas1-Cas2 and psDNA are provided to visualize the allosteric propagation patterns for all the different psDNA systems (original/psL/psH), at the half-site post-catalytic state (see **Figure 6** *top*). It becomes quite clear that the psH system is highly correlated overall, upon the first cleavage at PAMc, and the correlation or allosteric propagation proceeds largely via Cas2&2’ and dsDNA regions in the middle of complex. In contrast, psL appears to have the lowest correlation strength going through the psDNA, or reaching to the site2-non-PAMc region. In the original psDNA system, noticeable correlation still shows between the site2 and site1, along with a medium level of psDNA correlation propagation, comparing to psH and psL. Additionally, we have also calculated the Cas2&2’-dsDNA association energetics for respective Cas1-Cas2 psDNA complexes (in the half-site post-catalytic state). Notably, one can see that electrostatic interactions between the Cas2&2’ protein and the duplex part of DNA are strongest in the psH system (see **Figure 6** bottom), which again supports the idea that psDNA contributes directly to the correlation or allosteric communication in the Cas1-Cas2 acquisition complex by associating closely with Cas2&2’.

**Figure 6.**
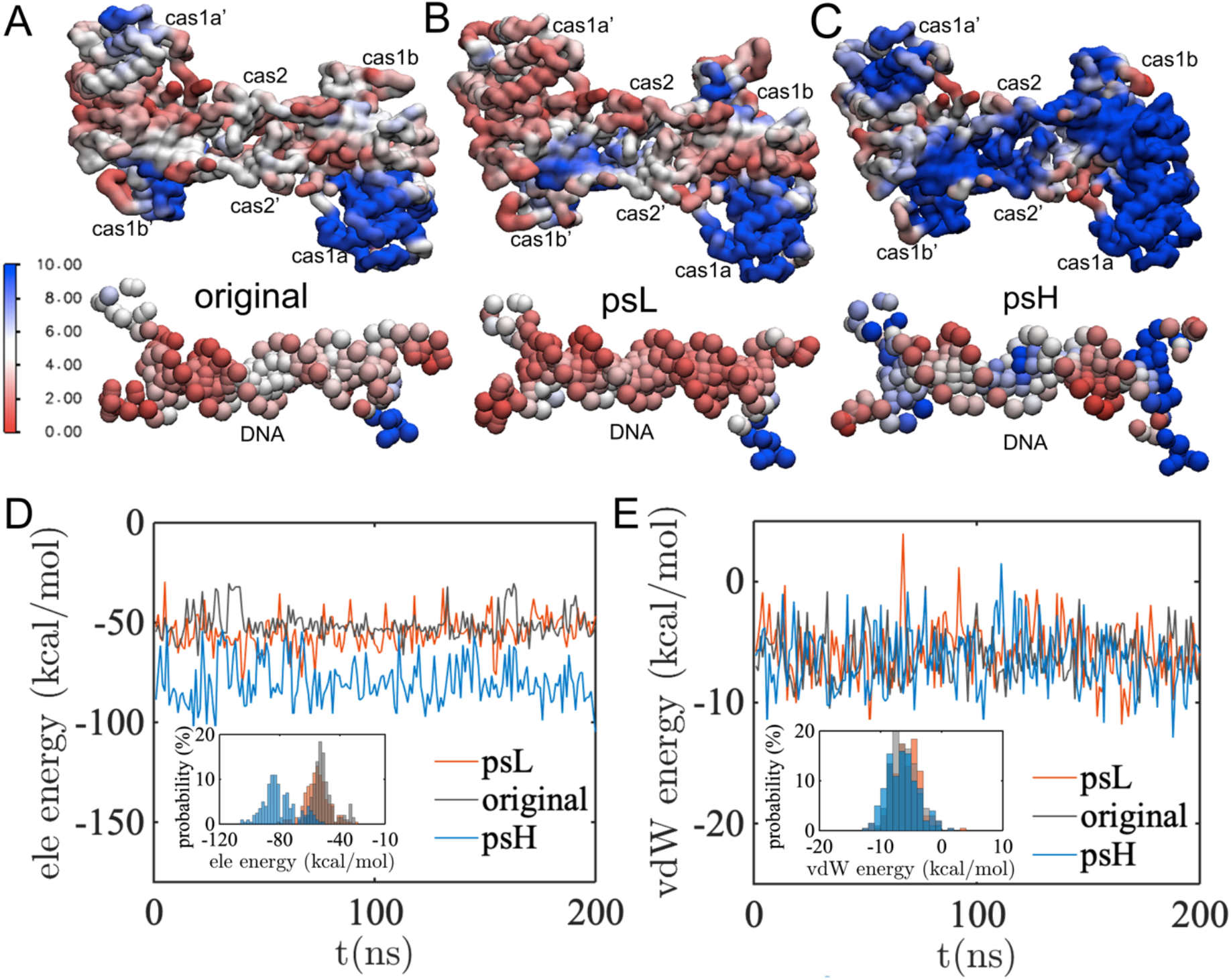
The color maps of correlation strength between the active site1 (bound with PAMc) and the rest of the protein-DNA complex viewed on the structures with different psDNA sequences, in the halfsite post-catalytic state. The correlation map on the Cas1-Cas2 structure and dual-fork psDNA (blue/white/red: high/medium/low correlation values), for the original psDNA sequences **(A)**, psL **(B)**, and psH **(C)**. Note the correlation data are the same as that shown in Figure 5, measured from the equilibrium MD simulations. **(D & E)** The electrostatic and vdW energies between Cas2&2’ and dsDNA for the various psDNA complexes: original/psL/psH (with average values -51±7, -55±8, and -80±11 kcal/mol for the electrostatic or ele energy, and -6.2±2.0, -5.7 ±2.4, and -6.4 ±2.5 kcal/mol for the vdW energy).

We also examined internal correlations for psL/psH and the original psDNA systems in the binding and the pre-catalytic state (see Supplementary **Figure S8**). Indeed, Cas1-Cas2 in complex with psH produces comparatively low correlation in the binding state, i.e., lower than that in the psL or the original system. In the pre-catalytic state (see detailed interaction characteristic and active site views in Supplementary **Figure S9 to S11**), however, the psH system correlation becomes slightly higher than that in the psL or the original DNA system. The correlation also seems to be mediated by Cas2&2’-DNA. The results suggest that the catalysis in the site1-PAMc is necessary to activate the allosteric propagation.

Last, we examined the HBs formed between Cas1-Cas2 and psDNA at the half-site post-catalytic state, comparing the systems with different psDNA sequences, i.e., the original/psL/psH. In particular, we calculated HB occupancies for these systems. For HB occupancies >50% between Cas1-Cas2 and DNA during the simulation, the psH system shows much more stabilized HBs than those in the original and psL systems (see **Figure 7** and Supplementary **Table S1**). One notices that there are three protein-DNA binding zones in the original psDNA system that show more HBs formed than that in the psL system: zone I (Cas1a&b & Cas2’ with nt 1-5 on the top strand of psDNA), zone II’ (Cas1b’ and Cas2&2’ with nt 13-16 on the top strand), and zone II (Cas1b and Cas2&2’ with nt 13-16 on the bottom strand). Furthermore, the regions show more HBs in the psH system than in the original system include an additional zone I’ (Cas1a’&b’ and Cas2 with nt 1-5 on the bottom strand), while zone II interaction is also enhanced in the psH system than in the original or psL system. As a result, one can see that Cas2&2’ dominate all the essential protein-DNA HB interaction zones connecting the two active sites, with enhanced HB interactions from the psL to the original and to the psH system. Hence, Cas2&2’ appear critical for the involved allosteric propagation.

**Figure 7.**
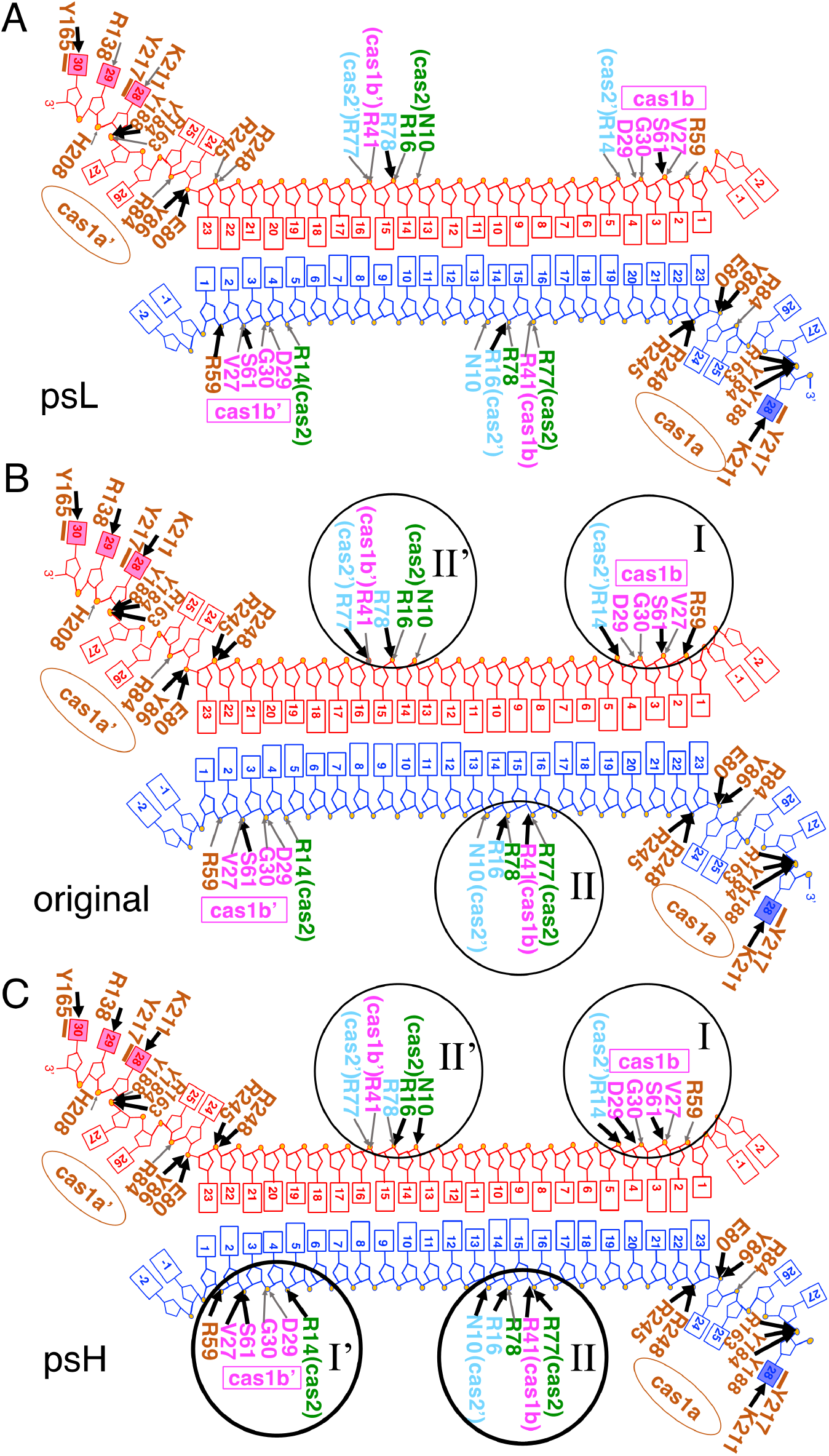
The schematics of the Cas1-Cas2 psDNA complex at the half-site post-catalytic state with high hydrogen-bonding (HB) occupancies obtained from three simulation systems with different psDNA sequences (psL /original /psH). **(A-C)** The HB occupancies are compared among the three simulation systems, and those HBs with occupancies >50% during the simulation time are labeled with black arrows, those <50% are labeled as well. Those seem to contribute to increase HB associations (from psL to original and to psH) are: N10, R14, R16, and R77 from Cas2, N10, R14 from Cas2’, D29 and R41 from Cas1b, R138, R163, K211, R245, and R248 from Cas1a’.

Among these HBs, R77, R78, R16 and R14 from Cas2&2’, interact with DNA substantially in the psH or in the original system, have also been found crucial experimentally. Recent high-resolution structural studies show that the mutants of R77 and R78 by Ala residues can reduce spacer acquisition efficiency; in addition, no spacer acquisition was observed for R16A and R14A mutants (27,28). The HB patterns at the protein-DNA interface and the protein-DNA interaction energies thus together suggest that the correlation builds up between the two active sites, primarily via the Cas2&2’-psDNA interaction region, and such allosteric communication becomes prominent in the psH system at the post-catalysis state, once the PAMc gets cleaved at site1, and the system awaits site2 to be engaged with the non-PAMc.

## DISCUSSION

In this work, we performed intensive atomistic simulations to investigate how CRISPR/Cas1-Cas2 binds, recognizes, and possibly conducts cleavages efficiently on the protospacer DNA or psDNA in the adaptation stage of CRISPR-Cas immunity process. To simulate Cas1-Cas2 consisting of Cas1a&b, Cas1a’&b’ and Cas2&2’ in stable association with psDNA, a large solvent box with a distance ~ 2 nm between the protein and boundary was utilized, which led to a large solvated simulation system with ~ 0.5 million atoms. A series of sub-microsecond equilibrium MD simulations were then performed to individual systems on i) the Cas1-Cas2 associating with psDNA of different PAMc/non-PAMc configurations at two active sites (dual PAMc, two single-PAMc, and dual-non-PAMc for binding), ii) the site1-PAMc & site2-non-PAMc configuration, systems from binding to pre-catalytic and to a half-site post-catalytic state (i.e., three states with the original psDNA), and iii) with two synthetic psDNA of low and high acquisition efficiency tested (psL and psH, two additional three-state systems), in a total of 12 simulations for several microseconds.

Interestingly, we found that even in association with two identical PAMc, the original two-fold symmetric Cas1-Cas2 demonstrate an asymmetric binding pattern between two active sites: the active site1 residues form stabilized HBs and stacking interactions with PAMc from one fork region of the psDNA, while the active site2 cannot bind stably or specifically to PAMc on the symmetric fork region. Therefore, it appears that only one active site of Cas1-Cas2 is capable of binding and recognition of PAMc at a time, which suits for locating only one PAMc on the psDNA for the Cas1-Cas2 target search. It then calls into question on how Cas1-Cas2 possibly binds and cleaves the non-PAMc sufficiently fast at site2, so that it allows formation of a half-site intermediate integration complex (29). In such a complex, the psDNA has its non-PAMc side of 3’-OH overhang linking to the CRISPR locus as if it is resulted from the first nucleophilic attack, which is likely only if the non-PAMc is cleaved almost as fast as the PAMc.

A solution to have non-PAMc on the psDNA cleaved sufficiently fast by Cas1-Cas2 is suggested from our simulations of Cas1-Cas2 complex with dual-forked psDNA, containing one PAMc and one non-PAMc at respective fork regions, i.e., in the site1-PAMc and site2-non-PAMc configuration. We found that even though the active site2 associates only loosely with non-PAMc initially (in the Cas1-Cas2 binding state with psDNA), once site-1 recognizes and cleaves on the associated PAMc (modeled as the half-site post-catalytic state), site-2 immediately becomes stabilized with the non-PAMc, so that to be ready for the second cleavage. The mechanism is summarized in schematics in **Figure 8**. The two active sites of Cas1-Cas2 can demonstrate negative cooperativity in a ‘seesaw’ manner, with one site being capable of binding specifically for the PAMc recognition, and conducting the first catalytic cleavage at that site; the other site catches up for tight DNA association to cleave nonspecifically, immediately after the first site cleavage, taking advantage of the negative cooperativity. Such negative or seesawing-type of cooperativity had been reported in other enzyme systems, for example, in the homo-dimeric insulin receptor, the membrane-spanning tyrosine kinase allowing insulin to dock into two binding pockets (53); in the Mo-bisPGD enzyme arsenite oxidase, which impacts on the early life metabolic reactions, with ‘the redox seesaw’ cooperativity induced by the pyranopterin ligands (54); and in the ‘seesaw’ model of enzyme regulation of mTORC1, in order to produce a nonlinear, ultrasensitive responses (55).

**Figure 8.**
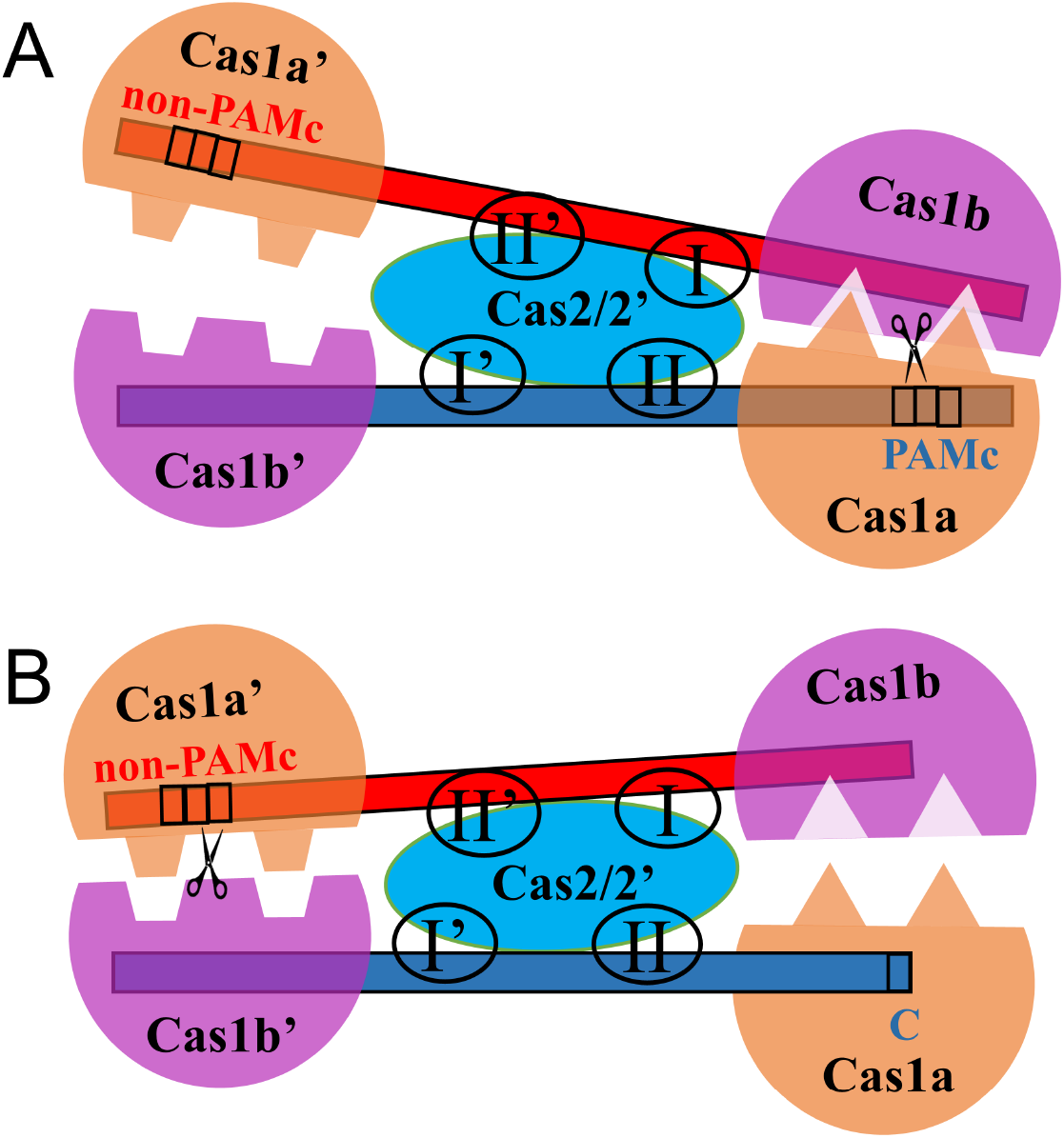
The schematics on the ‘see-saw’ type of negative cooperativity identified between the two active sites remotely (separated by ~ 10 nm) on the Cas1-Cas2 protospacer acquisition complex. The protospacer DNA is represented by red and blue strands. Cas1 and Cas2 consist of dimeric Cas1a&b, Cas1a’&b’ and Cas2&2’, which are colored differently and labeled. **(A)** The preferred binding to PAMc in the first active site of Cas1-Cas2 in psDNA binding and selection. **(B)** The consequent non-PAMc binding in the second active site of Cas1-Cas2, following immediately the PAMc cleavage at the first active site from (A). The Cas2&2’ interaction zones with the duplex psDNA are labeled (I&I’ and II&II’), where essential HBs formed. The interaction zones can play important roles for mediating allosteric propagation from the site1-PAMc to site2-non-PAMc.

The two active sites in the dimeric Cas1-Cas2 system are located symmetrically at two remote regions on the Cas1a&b and Cas1a’&b’, respectively, ~ 10 nm apart. The ~23 bp dsDNA region of the psDNA is bound primarily with Cas2&2’, in between Cas1a&b and Cas1a’&b’ containing the two active sites. In order to find out how the communication between the two remote sites achieves in the Cas1-Cas2 psDNA acquisition complex, we analyzed dynamic correlations within the complex from the MD trajectories, focusing on how the rest part of the protein-DNA complex correlates with the active site1 that is bound with PAMc, from the psDNA binding to the pre-catalytic state, and to the half-way post-catalytic state. Indeed, one could identify nontrivial correlation between the site2-non-PAMc, and site1-PAMc, and such correlation increases into the half-site post-catalytic state, right after the site1 cleavage on the PAMc. The correlation analysis, therefore, suggests that allosteric communication exists that propagates fluctuations at the active site1-PAMc cleavage to the remote site2-non-PAMc. More close examinations show that it is primarily the Cas2&2’-psDNA association, i.e, via the protein-DNA electrostatic interactions, as well as the hydrogen bonding interactions formed at the protein-DNA interface (particularly at the Cas2&2’-dsDNA interaction zones), that are responsible for such allosteric propagation. In such critical Cas2&2’-dsDNA interaction zones, R14, R16, R77, and R78 residues seem to play important roles to facilitate the allosteric propagation, and the mutants of these arginine residues reduce the protospacer acquisition efficiency significantly (27,28). Correspondingly, we propose that Cas2&2’-dsDNA embedded in the middle of the Cas1-Cas2 dimer mediates the two remotely located active site1 and site2, allosterically. With such mediation, negative cooperativity arises between the two remote sites, which allows the non-specific non-PAMc cleavages takes place quickly after the specific cleavage of PAMc by Cas1-Cas2.

To substantiate such a proposal, we further constructed Cas1-Cas2 complexes containing two synthetic DNA protospacers, psL and psH, which were experimentally identified with low and high integration efficiency, respectively (36). Similar MD simulations and correlation analyses were conducted to the modified Cas1-Cas2 psDNA complexes containing psL and psH, and the correlation patterns between site1-PAMc and site2-nonPAMc were monitored, in comparison with the original psDNA system. Notably, one finds that the inter-site correlation or allosteric propagation becomes significantly enhanced in the psH system, also into the half-site post-catalytic state, upon the PAMc cleavage at site1. Such correlation or allosteric propagation remains low, however, in the psL system. Hence, such studies further demonstrate that allosteric propagation from the site1-PAMc to the site2-non-PAMc can impact on the acquisition efficiency, and the allosteric effect can be particularly strong for certain psDNA sequences that tightly associate with Cas2&2’. Besides, in the allosteric propagation of Cas1-Cas2 via psDNA, a dsDNA ~ 23 bp (33 bp full length) actually sets a ruler length for the protospacer acquisition. It is interesting to notice that such a length is right about an upper bound (25-30 bp) for DNA allostery to be effective to regulate binding cooperativity of a pair of proteins at the neighboring locations on the DNA (56). Hence, such a psDNA length may indeed be optimized for supporting certain cooperativity between the two active sites on Cas1-Cas2.

Our work also suggests that Cas2&2’ plays a key role in the allosteric regulation of the protospacer DNA acquisition, besides its obvious structural role of bridging Cas1a&b and Cas1a’&b’ and in association with psDNA. Indeed, Cas2&2’ appears highly stable in the Cas1-Cas2 psDNA association complex in simulation, while psDNA is comparatively flexible. Consistently, Cas2&2’ and psDNA becomes more flexible into the half-site post-catalytic state than in the binding or pre-catalytic state. Such a role may also explain why Cas2 does not exhibit enzymatic activity in the acquisition process, even though it is enzymatically capable of making cleavages on single or doubled stranded DNA (16,26). Notably, recent studies discovered that two Cas1 dimer alone can still form a mini-integrase that binds psDNA at a shorter length (~18 bp), in the absence of Cas2 (57). Without Cas2, the two sites presumably come closer on the Cas1-psDNA complex (with a distance ~ 6 nm, shortened by 40% as that in the original Cas1-Cas2 system). It would be interesting to find out whether inter-site communication and cooperativity still exist in such a mini-integrase, and whether efficient protospacer acquisition is still a part of the ancestral Cas1 function, prior to Cas2 adoption (57). On the other hand, it is important to realize that Cas1-Cas2 protospacer acquisition process can also be supported by Cas3 (6), Cas4 (9), Cas9 and Casn2 in various CRISRP systems (58).

The PAM specific DNA acquisition of Cas1-Cas2 has actually been implemented for designing molecular recording (36,59). By using the nucleotide content, temporal ordering, and orientation of defined DNA sequences within a CRISPR array, Cas1-Cas2 seems to be able to encode arbitrary information within the genomes, and has a potential to record and store DNA information for long period of time. It was in such efforts of integrating synthetical DNA sequences via Cas1-Cas2, the psDNA sequences with particularly low and high acquisition efficiency (psL and psH) were identified (36). To allow Cas1-Cas2 to identify different PAMs in the synthetical approach, many mutants of Cas1-Cas2 have also been generated in the lab directed evolution (36). The physical mechanisms revealed in current work can be further tested, for example, in such a variety of Cas1-Cas2 mutants. It is expected that by combining information from experimental synthetical approaches, computational work would reveal substantial physical mechanisms to enable further rational molecular functional redesign.

## ACKNOWLEDGEMENT

We acknowledge the computational support from the Special Program for Applied Research on Super Computation of the NSFC Guangdong Joint Fund (the second phase) under Grant No. U1501501 and from the Beijing Computational Science Research Center (CSRC).

## REFERENCES

1. Barrangou, R., Fremaux, C., Deveau, H., Richards, M., Boyaval, P., Moineau, S., Romero, D.A. and Horvath, P. (2007) CRISPR Provides Acquired Resistance Against Viruses in Prokaryotes. Science, 315, 1709–1712.

2. Brouns, S.J.J., Jore, M.M., Lundgren, M., Westra, E.R., Slijkhuis, R., Snijders, A.P.L., Dickman, M.J., Makarova, K.S., Koonin, E.V. and Der Oost, J.V. (2008) Small CRISPR RNAs guide antiviral defense in prokaryotes. Science, 321, 960–964.

3. Sorek, R., Lawrence, C.M. and Wiedenheft, B. (2013) CRISPR-mediated adaptive immune systems in bacteria and archaea. Annual review of biochemistry, 82, 237–266.

4. Barrangou, R. and Marraffini, L.A. (2014) CRISPR-Cas systems: prokaryotes upgrade to adaptive immunity. Molecular cell, 54, 234–244.

5. Xiao, Y., Ng, S., Nam, K.H. and Ke, A. (2017) How type II CRISPR-Cas establish immunity through Cas1–Cas2-mediated spacer integration. Nature, 550, 137–141.

6. Fagerlund, R.D., Wilkinson, M.E., Klykov, O., Barendregt, A., Pearce, F.G., Kieper, S.N., Maxwell, H.W., Capolupo, A., Heck, A.J. and Krause, K.L. (2017) Spacer capture and integration by a type IF Cas1-Cas2–3 CRISPR adaptation complex. Proceedings of the National Academy of Sciences, 114, E5122–E5128.

7. Fineran, P.C. and Charpentier, E. (2012) Memory of viral infections by CRISPR-Cas adaptive immune systems: acquisition of new information. Virology, 434, 202–209.

8. McGinn, J. and Marraffini, L.A. (2019) Molecular mechanisms of CRISPR–Cas spacer acquisition. Nature Reviews Microbiology, 17, 7–12.

9. Lee, H., Dhingra, Y. and Sashital, D.G. (2019) The Cas4-Cas1-Cas2 complex mediates precise prespacer processing during CRISPR adaptation. Elife, 8, e44248.

10. Rollins, M.F., Chowdhury, S., Carter, J., Golden, S.M., Miettinen, H.M., Santiago-Frangos, A., Faith, D., Lawrence, C.M., Lander, G.C. and Wiedenheft, B. (2019) Structure reveals a mechanism of CRISPR-RNA-guided nuclease recruitment and anti-CRISPR viral mimicry. Molecular cell, 74, 132–142. e135.

11. Hale, C.R., Zhao, P., Olson, S., Duff, M.O., Graveley, B.R., Wells, L., Terns, R.M. and Terns, M.P. (2009) RNA-Guided RNA Cleavage by a CRISPR RNA-Cas Protein Complex. Cell, 139, 945–956.

12. Garneau, J.E., Dupuis, M.E., Villion, M., Romero, D.A., Barrangou, R., Boyaval, P., Fremaux, C., Horvath, P., Magadan, A.H. and Moineau, S. (2010) The CRISPR/Cas bacterial immune system cleaves bacteriophage and plasmid DNA. Nature, 468, 67–71.

13. Yosef, I., Goren, M.G. and Qimron, U. (2012) Proteins and DNA elements essential for the CRISPR adaptation process in Escherichia coli. Nucleic Acids Research, 40, 5569–5576.

14. Datsenko, K.A., Pougach, K., Tikhonov, A., Wanner, B.L., Severinov, K. and Semenova, E. (2012) Molecular memory of prior infections activates the CRISPR/Cas adaptive bacterial immunity system. Nature communications, 3, 1–7.

15. Swarts, D.C., Mosterd, C., Van Passel, M.W.J. and Brouns, S.J.J. (2012) CRISPR Interference Directs Strand Specific Spacer Acquisition. PLOS ONE, 7.

16. Nuñez, J.K., Kranzusch, P.J., Noeske, J., Wright, A.V., Davies, C.W. and Doudna, J.A. (2014) Cas1–Cas2 complex formation mediates spacer acquisition during CRISPR–Cas adaptive immunity. Nature structural & molecular biology, 21, 528.

17. Nunez, J.K., Lee, A.S., Engelman, A. and Doudna, J.A. (2015) Integrase-mediated spacer acquisition during CRISPR-Cas adaptive immunity. Nature, 519, 193–198.

18. Rollie, C., Schneider, S., Brinkmann, A.S., Bolt, E.L. and White, M.F. (2015) Intrinsic sequence specificity of the Cas1 integrase directs new spacer acquisition. eLife, 4.

19. Makarova, K.S., Wolf, Y.I., Alkhnbashi, O.S., Costa, F., Shah, S.A., Saunders, S.J., Barrangou, R., Brouns, S.J., Charpentier, E. and Haft, D.H. (2015) An updated evolutionary classification of CRISPR–Cas systems. Nature Reviews Microbiology, 13, 722–736.

20. Mohanraju, P., Makarova, K.S., Zetsche, B., Zhang, F., Koonin, E.V. and Der Oost, J.V. (2016) Diverse evolutionary roots and mechanistic variations of the CRISPR-Cas systems. Science, 353, 0.

21. Sternberg, S.H., Richter, H., Charpentier, E. and Qimron, U. (2016) Adaptation in CRISPR-Cas Systems. Molecular Cell, 61, 797–808.

22. Krupovic, M., Beguin, P. and Koonin, E.V. (2017) Casposons: mobile genetic elements that gave rise to the CRISPR-Cas adaptation machinery. Current Opinion in Microbiology, 38, 36–43.

23. Makarova, K.S., Wolf, Y.I., Iranzo, J., Shmakov, S., Alkhnbashi, O.S., Brouns, S.J.J., Charpentier, E., Cheng, D.R., Haft, D.H. and Horvath, P. (2020) Evolutionary classification of CRISPR–Cas systems: a burst of class 2 and derived variants. Nature Reviews Microbiology, 18, 67–83.

24. Beloglazova, N., Brown, G., Zimmerman, M.D., Proudfoot, M., Makarova, K.S., Kudritska, M., Kochinyan, S., Wang, S., Chruszcz, M. and Minor, W. (2008) A Novel Family of Sequence-specific Endoribonucleases Associated with the Clustered Regularly Interspaced Short Palindromic Repeats. Journal of Biological Chemistry, 283, 20361–20371.

25. Babu, M., Beloglazova, N., Flick, R., Graham, C., Skarina, T., Nocek, B., Gagarinova, A., Pogoutse, O., Brown, G. and Binkowski, A. (2011) A dual function of the CRISPR–Cas system in bacterial antivirus immunity and DNA repair. Molecular Microbiology, 79, 484–502.

26. Nam, K.H., Ding, F., Haitjema, C.H., Huang, Q., Delisa, M.P. and Ke, A. (2012) Double-stranded endonuclease activity in Bacillus halodurans clustered regularly interspaced short palindromic repeats (CRISPR)-associated Cas2 protein. Journal of Biological Chemistry, 287, 35943–35952.

27. Wang, J., Li, J., Zhao, H., Sheng, G., Wang, M., Yin, M. and Wang, Y. (2015) Structural and Mechanistic Basis of PAM-Dependent Spacer Acquisition in CRISPR-Cas Systems. Cell, 163, 840–853.

28. Nunez, J.K., Harrington, L.B., Kranzusch, P.J., Engelman, A. and Doudna, J.A. (2015) Foreign DNA capture during CRISPR–Cas adaptive immunity. Nature, 527, 535–538.

29. Wright, A.V., Liu, J.-J., Knott, G.J., Doxzen, K.W., Nogales, E. and Doudna, J.A. (2017) Structures of the CRISPR genome integration complex. Science, 357, 1113–1118.

30. Xiao, Y., Budhathoki, J., Ng, S., Nam, K.H. and Ke, A. (2018) Spacer acquisition mechanism in type II-A CRISPR system.

31. Budhathoki, J.B., Xiao, Y., Schuler, G., Hu, C., Cheng, A., Ding, F. and Ke, A. (2020) Real-time observation of CRISPR spacer acquisition by Cas1–Cas2 integrase. Nature Structural & Molecular Biology, 27, 489–499.

32. Horvath, P., Romero, D.A., Coutemonvoisin, A., Richards, M., Deveau, H., Moineau, S., Boyaval, P., Fremaux, C. and Barrangou, R. (2008) Diversity, activity, and evolution of CRISPR loci in Streptococcus thermophilus. Journal of Bacteriology, 190, 1401–1412.

33. Goren, M.G., Yosef, I., Auster, O. and Qimron, U. (2012) Experimental Definition of a Clustered Regularly Interspaced Short Palindromic Duplicon in Escherichia coli. Journal of Molecular Biology, 423, 14–16.

34. Deveau, H., Barrangou, R., Garneau, J.E., Labonté, J., Fremaux, C., Boyaval, P., Romero, D.A., Horvath, P. and Moineau, S. (2008) Phage response to CRISPR-encoded resistance in Streptococcus thermophilus. Journal of bacteriology, 190, 1390–1400.

35. Mojica, F.J., Díez-Villaseñor, C., García-Martínez, J. and Almendros, C. (2009) Short motif sequences determine the targets of the prokaryotic CRISPR defence system. Microbiology, 155, 733–740.

36. Shipman, S.L., Nivala, J., Macklis, J.D. and Church, G.M. (2016) Molecular recordings by directed CRISPR spacer acquisition. Science, 353.

37. Price, D.J. and Brooks, C.L. (2004) A modified TIP3P water potential for simulation with Ewald summation. Journal of Chemical Physics, 121, 10096–10103.

38. Mackerell, A.D., Bashford, D., Bellott, M., Dunbrack, R.L., Evanseck, J.D., Field, M.J., Fischer, S., Gao, J., Guo, H. and Ha, S. (1998) All-Atom Empirical Potential for Molecular Modeling and Dynamics Studies of Proteins †. Journal of Physical Chemistry B, 102, 3586–3616.

39. Berendsen, H.J.C., Der Spoel, D.V. and Van Drunen, R. (1995) GROMACS - A MESSAGEPASSING PARALLEL MOLECULAR-DYNAMICS IMPLEMENTATION. Computer Physics Communications, 91, 43–56.

40. Abraham, M., van der Spoel, D., Lindahl, E. and Hess, B. (2016) GROMACS User Manual, version 5.1.2; GROMACS Development Team. http://manual.gromacs.org/5.1.2/ReleaseNotes/index.html.

41. Joung, I. and Cheatham, T.E. (2008) Determination of Alkali and Halide Monovalent Ion Parameters for Use in Explicitly Solvated Biomolecular Simulations. Journal of Physical Chemistry B, 112, 9020–9041.

42. Joung, I. and Cheatham, T.E. (2009) Molecular Dynamics Simulations of the Dynamic and Energetic Properties of Alkali and Halide Ions Using Water-Model-Specific Ion Parameters. Journal of Physical Chemistry B, 113, 13279–13290.

43. Lindorfflarsen, K., Piana, S., Palmo, K., Maragakis, P., Klepeis, J.L., Dror, R.O. and Shaw, D.E. (2010) Improved side - chain torsion potentials for the Amber ff99SB protein force field. Proteins, 78, 1950–1958.

44. Guy, A.T., Piggot, T.J. and Khalid, S. (2012) Single-Stranded DNA within Nanopores: Conformational Dynamics and Implications for Sequencing; a Molecular Dynamics Simulation Study. Biophysical Journal, 103, 1028–1036.

45. Darden, T., York, D.M. and Pedersen, L.G. (1993) Particle mesh Ewald: An N · log(N) method for Ewald sums in large systems. Journal of Chemical Physics, 98, 10089–10092.

46. Essmann, U., Perera, L., Berkowitz, M.L., Darden, T., Lee, H. and Pedersen, L.G. (1995) A smooth particle mesh Ewald method. Journal of Chemical Physics, 103, 8577–8593.

47. Parrinello, M. and Rahman, A. (1981) Polymorphic transitions in single crystals: A new molecular dynamics method. Journal of Applied Physics, 52, 7182–7190.

48. Bussi, G., Donadio, D. and Parrinello, M. (2007) Canonical sampling through velocity rescaling. Journal of Chemical Physics, 126, 014101.

49. Nosé, S. and Klein, M. (1983) Constant pressure molecular dynamics for molecular systems. Molecular Physics, 50, 1055–1076.

50. Li, S., Olson, W.K. and Lu, X. (2019) Web 3DNA 2.0 for the analysis, visualization, and modeling of 3D nucleic acid structures. Nucleic Acids Research, 47.

51. Yu, J., Ha, T. and Schulten, K. (2007) How directional translocation is regulated in a DNA helicase motor. Biophysical journal, 93, 3783–3797.

52. Dai, L., Flechsig, H. and Yu, J. (2017) Deciphering intrinsic inter-subunit couplings that lead to sequential hydrolysis of F1-ATPase ring. Biophysical Journal, 113, 1440–1453.

53. Vashisth, H. and Abrams, C.F. (2010) Docking of insulin to a structurally equilibrated insulin receptor ectodomain. Proteins, 78, 1531–1543.

54. Duval, S., Santini, J.M., Lemaire, D., Chaspoul, F., Russell, M.J., Grimaldi, S., Nitschke, W. and Schoeppcothenet, B. (2016) The H-bond network surrounding the pyranopterins modulates redox cooperativity in the molybdenum-bisPGD cofactor in arsenite oxidase. Biochimica et Biophysica Acta, 1857, 1353–1362.

55. Rahman, A. and Haugh, J.M. (2017) Kinetic Modeling and Analysis of the Akt/Mechanistic Target of Rapamycin Complex 1 (mTORC1) Signaling Axis Reveals Cooperative, Feedforward Regulation. Journal of Biological Chemistry, 292, 2866–2872.

56. Kim, S., Broströmer, E., Xing, D., Jin, J., Chong, S., Ge, H., Wang, S., Gu, C., Yang, L. and Gao, Y.Q. (2013) Probing allostery through DNA. Science, 339, 816–819.

57. Wright, A.V., Wang, J.Y., Burstein, D., Harrington, L.B., Paez-Espino, D., Kyrpides, N.C., Iavarone, A.T., Banfield, J.F. and Doudna, J.A. (2019) A functional mini-integrase in a two-protein type VC CRISPR system. Molecular cell, 73, 727–737. e723.

58. Wilkinson, M., Drabavicius, G., Silanskas, A., Gasiunas, G., Siksnys, V. and Wigley, D.B. (2019) Structure of the DNA-bound spacer capture complex of a type II CRISPR-Cas system. Molecular cell, 75, 90–101. e105.

59. Shipman, S.L., Nivala, J., Macklis, J.D. and Church, G.M. (2017) CRISPR–Cas encoding of a digital movie into the genomes of a population of living bacteria. Nature, 547, 345–349.

